# *In silico* secretome characterization of clinical *Mycobacterium abscessus* isolates provides insights into antigenic differences

**DOI:** 10.1101/2020.10.22.349720

**Authors:** Fernanda Cornejo-Granados, Thomas A. Kohl, Flor Vásquez Sotomayor, Sönke Andres, Rogelio Hernández-Pando, Juan Manuel Hurtado-Ramírez, Christian Utpatel, Stefan Niemann, Florian P. Maurer, Adrian Ochoa-Leyva

## Abstract

*Mycobacterium abscessus* (MAB) is a widely disseminated pathogenic non-tuberculous mycobacterium (NTM). Like with *M. tuberculosis* complex (MTBC), excreted / secreted (ES) proteins play an essential role for its virulence and survival inside the host. ES proteins contain highly immunogenic proteins, which are of interest for novel diagnostic assays and vaccines. Here, we used a robust bioinformatics pipeline to predict the secretome of the *M. abscessus* ATCC 19977 reference strain and fifteen clinical isolates belonging to all three MAB subspecies, *M. abscessus* subsp. *abscessus*, *M. abscessus* subsp. *bolletii,* and *M. abscessus* subsp. *massiliense.* We found that ~18% of the proteins encoded in the MAB genomes were predicted as secreted and that the three MAB subspecies shared > 85 % of the predicted secretomes. MAB isolates with a rough (R) colony morphotype showed larger predicted secretomes than isolates with a smooth (S) morphotype. Additionally, proteins exclusive to the secretomes of MAB R variants had higher antigenic densities than those exclusive to S variants, independently of the subspecies. For all investigated isolates, ES proteins had a significantly higher antigenic density than non-ES proteins. We identified 337 MAB ES proteins with homologues in previously investigated *M. tuberculosis* secretomes. Among these, 222 have previous experimental support of secretion, and some proteins showed homology with protein drug targets reported in the DrugBank database. The predicted MAB secretomes showed a higher abundance of proteins related to quorum-sensing and Mce domains as compared to MTBC indicating the importance of these pathways for MAB pathogenicity and virulence. Comparison of the predicted secretome of *M. abscessus* ATCC 19977 with the list of essential genes revealed that 99 secreted proteins corresponded to essential proteins required for *in vitro* growth. All predicted secretomes were deposited in the Secret-AAR web-server (http://microbiomics.ibt.unam.mx/tools/aar/index.php).

## Introduction

Non-tuberculous mycobacteria (NTM) are widely disseminated, mostly saprophytic and partly opportunistic bacteria. The prevalence of NTM in clinical specimens has increased globally, and in some industrialized countries, infections caused by NTM are becoming more common than tuberculosis (TB). Infections caused by *M. abscessus* (MAB) are particularly challenging to manage due to the extensive innate resistance of MAB against a wide spectrum of clinically available antimicrobials (Nessar et al., 2012). MAB causes mostly pulmonary and occasionally extrapulmonary infections that can affect all organs in the human body (Lee et al., 2015). Current treatments for MAB induced pulmonary disease are long, associated with severe side effects and a cure rate below 50 % (Chen et al., 2019; Jarand et al., 2011; Sanguinetti et al., 2001). MAB is comprised of three subspecies, *M. abscessus* subsp. *abscessus, M. abscessus* subsp. *bolletii* and *M. abscessus* subsp. *massiliense,* hereafter referred to as MABA, MAB_B_, and MAB_M_, respectively (Tortoli et al., 2016). MAB isolates can show smooth (S) and rough (R) colony morphotypes, a trait that relies on the presence (S) or absence (R) of surface-associated glycopeptidolipids (GPLs) and that correlates with the virulence of the strain (Abeles and Pride, 2014; Howard et al., 2006; Ripoll et al., 2007) (Gutiérrez et al., 2018)). Transitioning from high-GPL to low-GPL production is observed in sequential MAB isolates obtained from patients with chronic underlying pulmonary disease. In these patients, S-to-R conversion is thought to present a selective advantage as the aggregative properties of MAB R variants strongly affect intracellular survival. In addition, a propensity to grow as extracellular cords allows these low-GPL producing bacilli to escape innate immune defenses (Gutiérrez et al., 2018).

The complete set of proteins excreted / secreted (ES) by a bacterial cell is referred to as its secretome. The secretome is involved in critical biological processes such as cell adhesion, migration, cell-to-cell communication and signal transduction (Tjalsma et al., 2004) ES proteins are considered an important source of molecules for serological diagnosis. Also, secreted proteins can be highly antigenic due to their immediate availability to the host immune system and are thus of interest in vaccinology (Daugelat et al., 1992; Zheng et al., 2013). So far, there have been few efforts to experimentally determine the secretome of MAB, and in particular, the secretomes of clinical MAB isolates (Gupta et al., 2009; Laencina et al., 2018; Shin et al., 2010; Yadav and Gupta, 2012). Nowadays, sequencing and bioinformatics strategies can be explored for the systematized prediction of ES proteins from bacterial genomes (Cornejo-Granados et al., 2017; Gomez et al., 2015). Recently, a robust bioinformatics pipeline for predicting and analyzing the complete *in silico* secretome of two clinical *M. tuberculosis* (MTB) genomes was published in 2017 by our group showing higher overall agreement with an experimental secretome compiled from literature than two previously reported secretomes for *M. tuberculosis* H37Rv (Cornejo-Granados et al., 2017).

To gain further insights into MAB ES proteins and their association with virulence and pathogenicity we sequenced and assembled the genomes of fifteen clinical MAB isolates belonging to all three subspecies including S and R morphotypes. We then adapted the bioinformatics strategy previously established for MTB to predict and analyze the complete set of ES proteins encoded in these isolates and in the *M. abscessus* ATCC 19977 type strain, and compared it with our previous findings for MTB (Cornejo-Granados et al., 2017).

## Materials and methods

### Ethics statement

Ethical review and approval was not required for the study as all work was performed on bacterial isolates archived at the strain repository of the National Reference Center for Mycobacteria in Borstel, Germany, in accordance with local legislation and institutional requirements. In particular, no data allowing identification of the affected patients was shared or released and no human DNA was sequenced or analyzed.

### Clinical isolates

We selected fifteen MAB clinical isolates comprising members of all MAB subspecies (MABA, n = 7; MABB, n = 4; MABM, n = 4) and both S (n = 8) and R (n = 6) morphotypes (not determined, n = 1). The strains were isolated from different biological sources representing both pulmonary colonization / infection (sputum, n = 10) and extrapulmonary samples (skin, n = 1; soft tissue, n = 1; lymph nodes, n = 2; blood, n = 1) (Table 1 and S1). For routine diagnostic purposes, species identification was performed using GenoType NTM-DR line probe assays (HAIN Lifescience, Nehren, Germany) and sequencing of the 16S and *rpoB* genes as described previously (Adekambi et al., 2003).

**Table 1.**
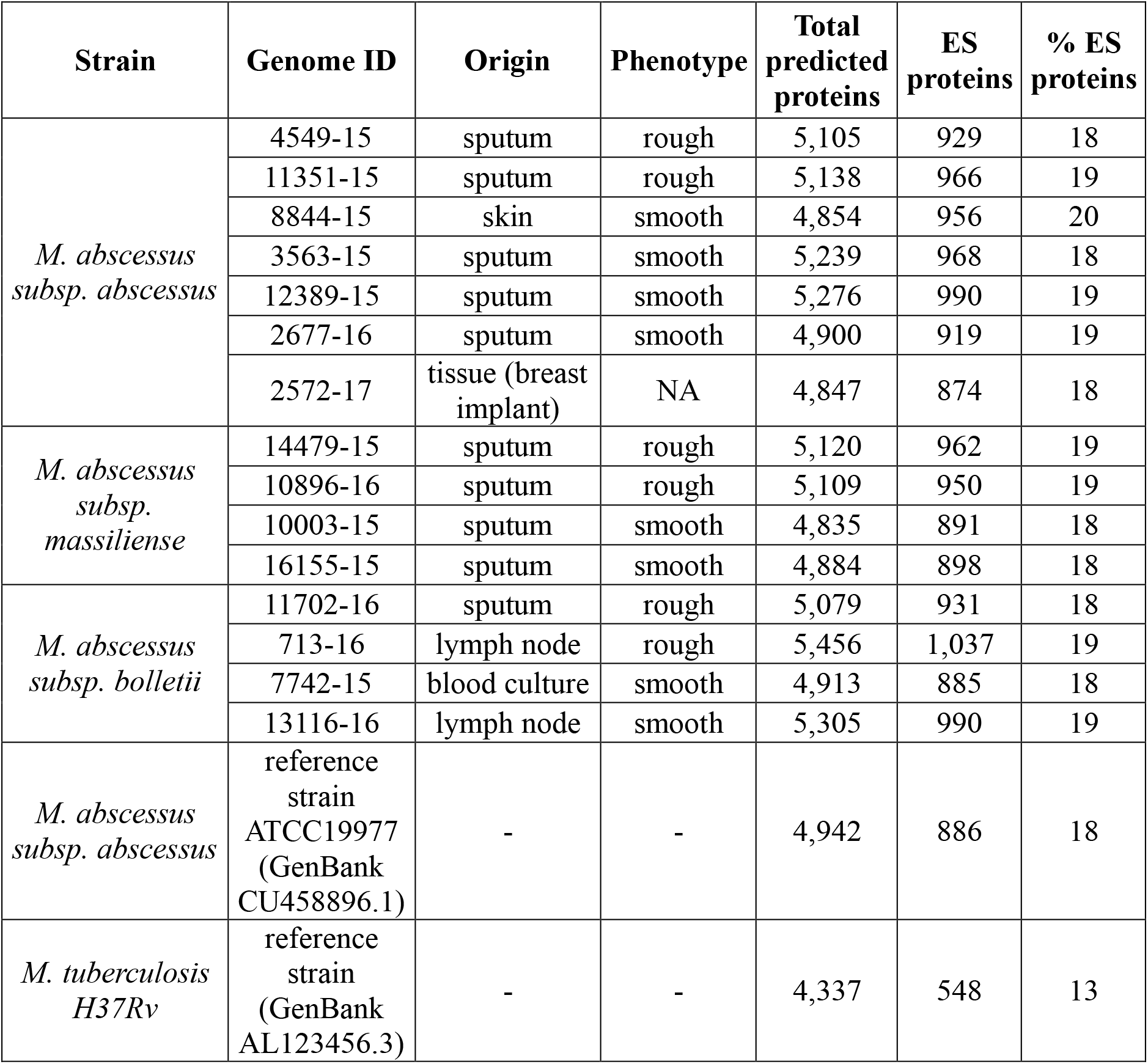
Clinical isolates metadata and number of ES proteins.

### Whole genome sequencing and genome assemblies

Genomic DNA (gDNA) of the 15 MAB clinical isolates was extracted from solid cultures using a Centrimonium bromide chloroform DNA extraction protocol as previously described (De Almeida et al., 2013). DNA libraries were constructed with the Nextera XT kit from Illumina and sequenced on the Illumina MiSeq benchtop platform with a v3 chemistry paired-end run and a read lenght of 2×300 bp. We processed the resulting reads with Trimmomatic (Bolger et al., 2014), clipping the Illumina adapter sequences and trimming the reads with a sliding window of 20 bp looking for quality >30 and discarding all reads shorter than 100 bp. Trimmed reads were used to construct *de novo* assemblies using SPADES (Nurk et al., 2013) with default parameters and the --careful option enabled. Then, each assembly was analyzed with RAST (Aziz et al., 2008) to obtain all the open reading frames (ORFs). Additionally, we predicted the ORFs from the deposited genome sequence of the *M. abscessus* ATCC 19977 type strain (GenBank CU458896.1) (Supplementary Table S1).

### Secretome prediction

The complete set of predicted ORFs was independently analyzed for each genome using the bioinformatics pipeline previously reported by Cornejo-Granados et al., 2017 and summarized in Supplementary Fig. S1. Briefly, we used six different feature-based tools (SignalP, SecretomeP, LipoP, TatP, TMHMM and Phobius) (Bendtsen et al., 2005a; 2005b; Petersen et al., 2011; Sonnhammer et al., 1998) (Juncker et al., 2003; Käll et al., 2007) to identify ES proteins by the different secretion pathways and to remove the ones that had transmembrane domains (Supplementary Fig. S1). The proteins assigned as not-secreted (non-ES) were further classified into transmembrane proteins (TM) if they showed the presence of transmembrane domains with TMHMM 2.0 (Sonnhammer et al., 1998), and into intracellular proteins (incell) if they did not contain any transmembrane domains.

### Annotation and comparative analysis of secreted proteins

To assign functional annotations to the proteins present in our genomes, we performed a BLASTP query of those proteins against the non-redundant (nr) complete database using Blast2GO (Conesa and Götz, 2008) with an E-value cut-off set at 1.0E-3. Furthermore, all proteins were associated with protein families through InterProScan (Zdobnov and Apweiler, 2001) and functionally mapped to Gene Ontology (GO) terms by setting the following parameters: E-value-hi-filter: 1.0E-3; Annotation cut-off: 55; GO weight: 5 and Hsp-Hit Coverage cut-off: 0. Blast2GO was then used to identify over- and under-represented GO and Enzyme Commission (EC) numbers in the ES proteins by setting the significance filter p-value to ≤ 0.05. Also, we used the KEGG Automatic Annotation Server (KAAS) database (Moriya et al., 2007) to assign the pathway annotation to the secreted proteins using the BBH (bidirectional best hit) method and the gene data set assigned to *Mycobacterium.*

To determine differences between the predicted secretomes in relation to MAB subspecies and morphotype, we established core secretomes by performing a bidirectional best-hit BLASTP search (E-value 1.0E-3) between the ES proteins of all genomes belonging to the respective subspecies and morphotypes. Then, we identified the shared and unique proteins for each comparison. Additionally, we determined the ES proteins shared between the MAB reference strain ATCC 19977 and *M. tuberculosis* H37Rv predicted and experimental secretomes (Cornejo-Granados et al., 2017). The resulting shared proteins were further investigated for sequence similarities against known drug targets available on the Drug Bank database (http://www.drugbank.ca/), setting the E-value to 1.0E-3 and all other options to default. In Supplementary Table S2, we show all proteins that have similarity with an approved drug target, as well as the drugs that can affect said target.

Additionally, we analyzed the presence of the core secretomes in twenty *M. abscessus* genomes per subspecies downloaded from NCBI (Supplementary Table S3). To this end, each downloaded genome was analyzed with RAST to obtain all the open reading frames (ORFs). Next, we performed a BLASTP search (E-value 1.0E-3) of each core secretome against each genome of the corresponding subspecies, and all hit proteins were considered homologs.

### Calculation of the Abundance of Antigenic Regions

The Abundance of Antigenic Regions (AAR) value is used to estimate the antigenic density of a protein by calculating the number of antigenic regions and normalizing it to the sequence length (Gomez et al., 2015). Of note, proteins with higher antigenic densities have lower AAR values. For this study, we calculated the AAR value for each protein in each data set using the Secret-AAR web-server (http://microbiomics.ibt.unam.mx/tools/aar/index.php) and reported the average unless stated otherwise (Cornejo-Granados et al., 2018). Then, we used a Mann-Whitney statistical test to establish any significant differences between the AAR values of the different protein data sets.

### Data availability

The reference genomes analyzed for *M. abscessus* ATCC19977 and *M. tuberculosis* H37Rv were taken from NCBI, under GenBank IDs CU458896.1 and NC_000962.3, respectively. The Whole Genome Shotgun project has been deposited at NCBI, under BioProject PRJNA646278. All the predicted secretomes were deposited in the Secret-AAR web-server (http://microbiomics.ibt.unam.mx/tools/aar/index.php).

## Results

### Genome assembly, secretome prediction and annotation

We sequenced the genomes of fifteen pulmonary and extrapulmonary (skin, tissue, and lymph node, blood) MAB isolates obtained from patients in Germany comprising all three MAB subspecies (Table 1 and S1). For each genome, we obtained an average of 2,601,444 quality-filtered reads. After *de novo* assembly, we obtained from 38 to 78 contigs (mean = 58 contigs) with genome coverage of 217- to 368-fold (mean = 310-fold) and with an average of 5,082 total proteins per genome (Supplementary Table S4).

We used a bioinformatics pipeline previously reported by our group (Cornejo-Granados et al., 2017) to predict the full secretome of all MAB clinical isolates and the widely used reference strain *M. abscessus* ATCC 19977 (GenBank CU458896.1) (Supplementary Fig. S1). We obtained an average of 939 ES proteins per genome, representing ~ 18% of the total proteome (Table 1). The predicted secretome for the MAB reference strain consisted of 886 proteins. From these, all the proteins showed a BLASTP hit against the NR database, and only 494 (55.8%) could be annotated with GO terms.

We analyzed the over-representation of GO terms in the secretome of *M. abscessus* ATCC 19977 as compared to the whole genome. The most significantly enriched GO-terms were: “lytic vacuole” (p = 9.37E-04) and “fungal-type vacuole” (p = 0.004) in Cellular Component (Fig. 1A), “serine-type carboxypeptidase” (p = 1.83E-04), and “serine-type D-Ala-D-Ala carboxypeptidase” (p = 1.83E-04) activities in Molecular Function (Fig. 1B) and “response to inorganic substance” (p = 5.68E-04) and “cellular response to oxygen radical” (p = 0.001) in the Biological Process category (Fig. 1C). The KEGG pathway mapping of the ES proteins showed that 214 proteins (24.2 %) could be assigned to 100 different KEGG pathways (Table 2), with the ABC transporter pathway being the most abundant (n= 13, 1.47 %). Additionally, serine-type D-Ala-D-Ala carboxypeptidases (p = 1.83E-04) and peptidases (p = 8.40E-04) were the most significantly abundant enzymes according to the Enzyme Commission (EC) Classes (Figure S2), while the Mce/MiaD and PknH-like extracellular domains were the most enriched protein domains (Table 3). Of note, comparably few sequences were assigned to the PE/PPE category (n = 3). Notably, after comparing the predicted secretome of *M. abscessus* ATCC 19977 with a list of essential genes published by (Laencina et al., 2018), we found that 99 (11.17 %) of the predicted ES proteins, corresponded to essential proteins required for *in vitro* growth.

**Figure 1.**
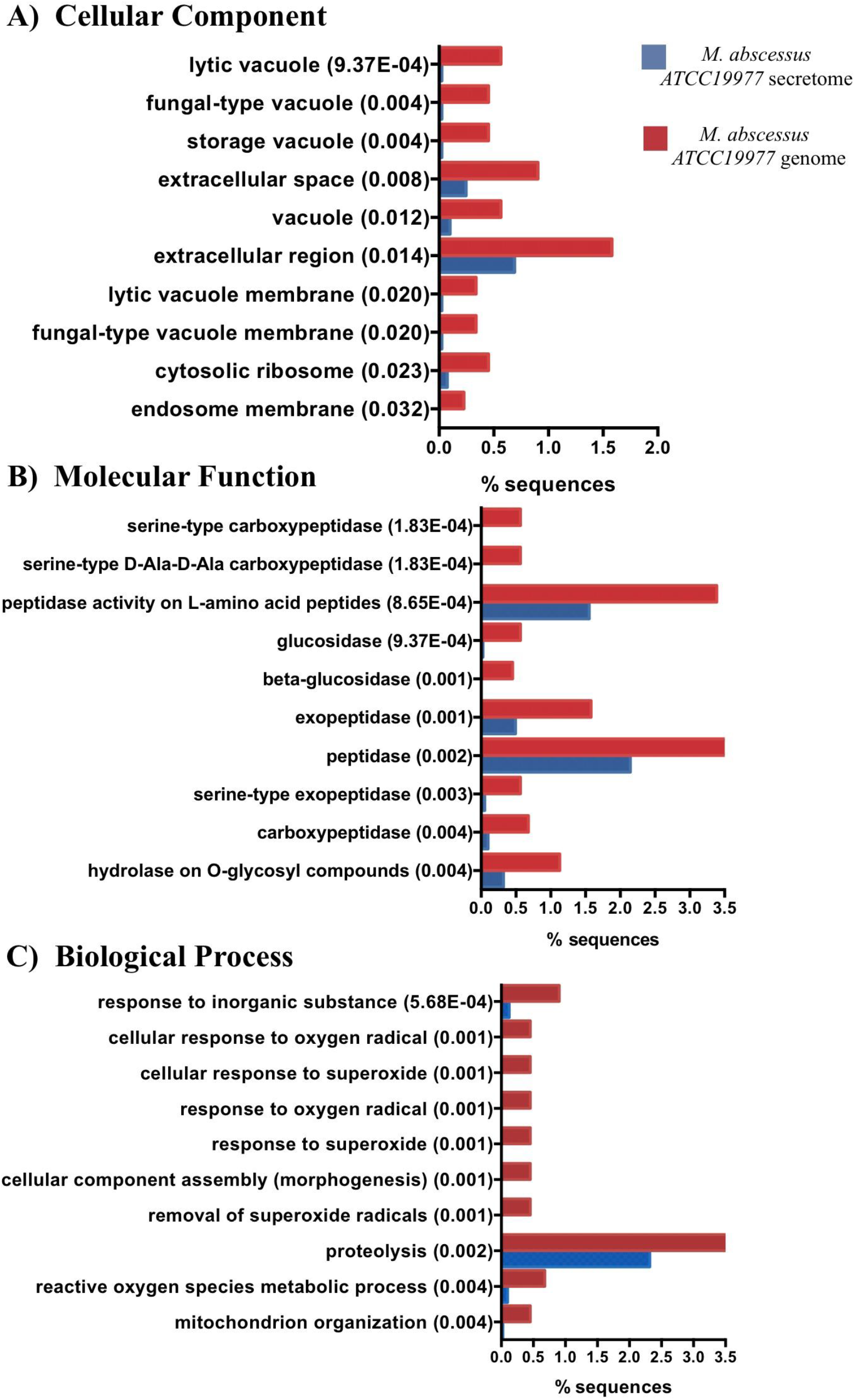
GO enrichment analysis for the *M. abscessus* ATCC 19977 reference strain. Top 10 most enriched GO terms for the *M. abscessus* ATCC 19977 secretome (blue) and complete genome (red) in three categories: A) Cellular Component, B) Molecular Function and C) Biological Process.

**Table 2.** Top 10 KEGG pathways assigned for*M. abscessus* ATCC19977 ES proteins.

| Ranking | Pathway name | Number of represented ES proteins (%) |
| --- | --- | --- |
| 1 | ABC transporters | 13 (1.47) |
| 2 | Two-component system | 9 (1.02) |
| 3 | Quorum sensing | 6 (0.68) |
| 4 | Oxidative phosphorylation | 4 (0.45) |
| 5 | Sulfur metabolism | 4 (0.45) |
| 6 | Glycerolipid metabolism | 4 (0.45) |
| 7 | Peptidoglycan biosynthesis | 4 (0.45) |
| 8 | Protein export | 4 (0.45) |
| 9 | Starch and sucrose metabolism | 3 (0.34) |
| 10 | Glyoxylate and dicarboxylate metabolism | 3 (0.34) |

**Table 3.**
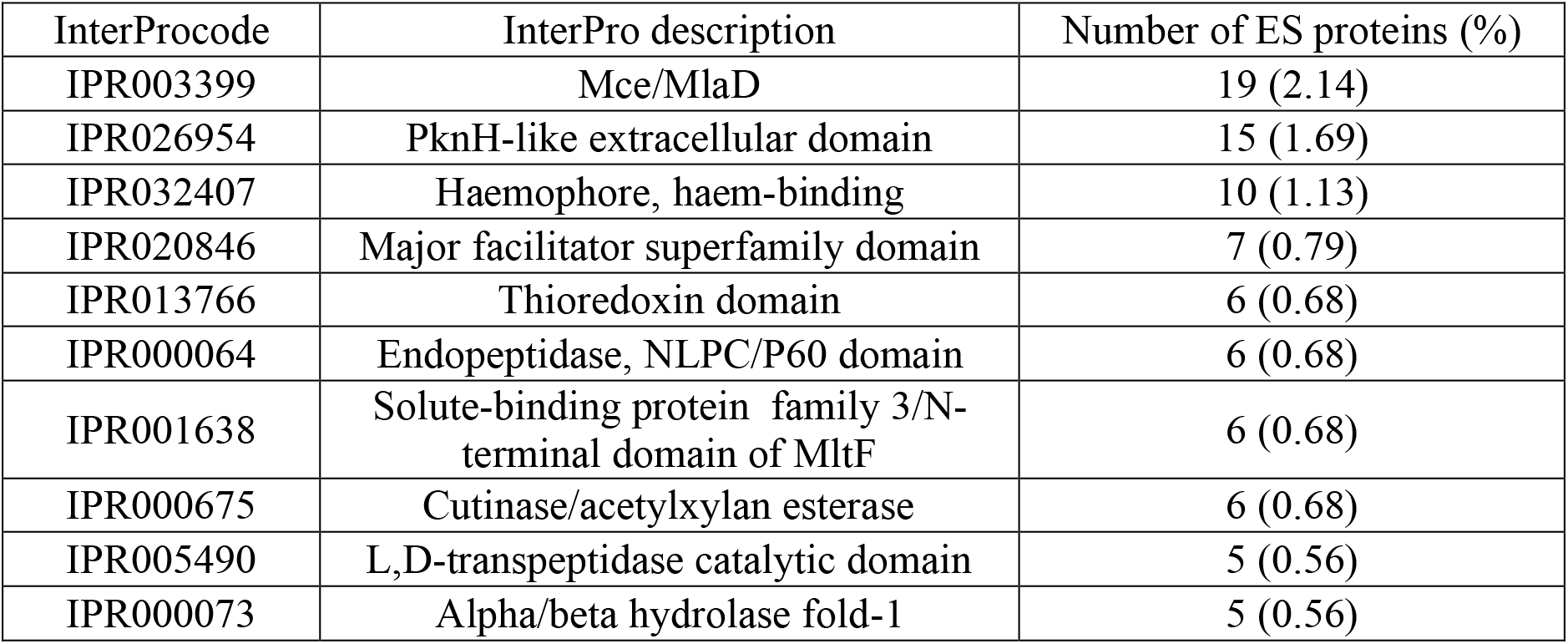
Top 10 most represented protein domains in *M. abscessus* ATCC19977 secretome.

### Comparison of *M. abscessus* subspecies core secretomes

We analyzed the differences between the predicted secretomes of the three MAB subspecies. To this end, we defined the core secretome of each subspecies as the set of proteins shared between all secretomes of isolates belonging to MABA, MABB, and MABM, respectively. The resulting core secretomes contained 735 (MABA), 794 (MAB_B_), and 813 (MABM) proteins (Fig 2A).

**Figure 2.**
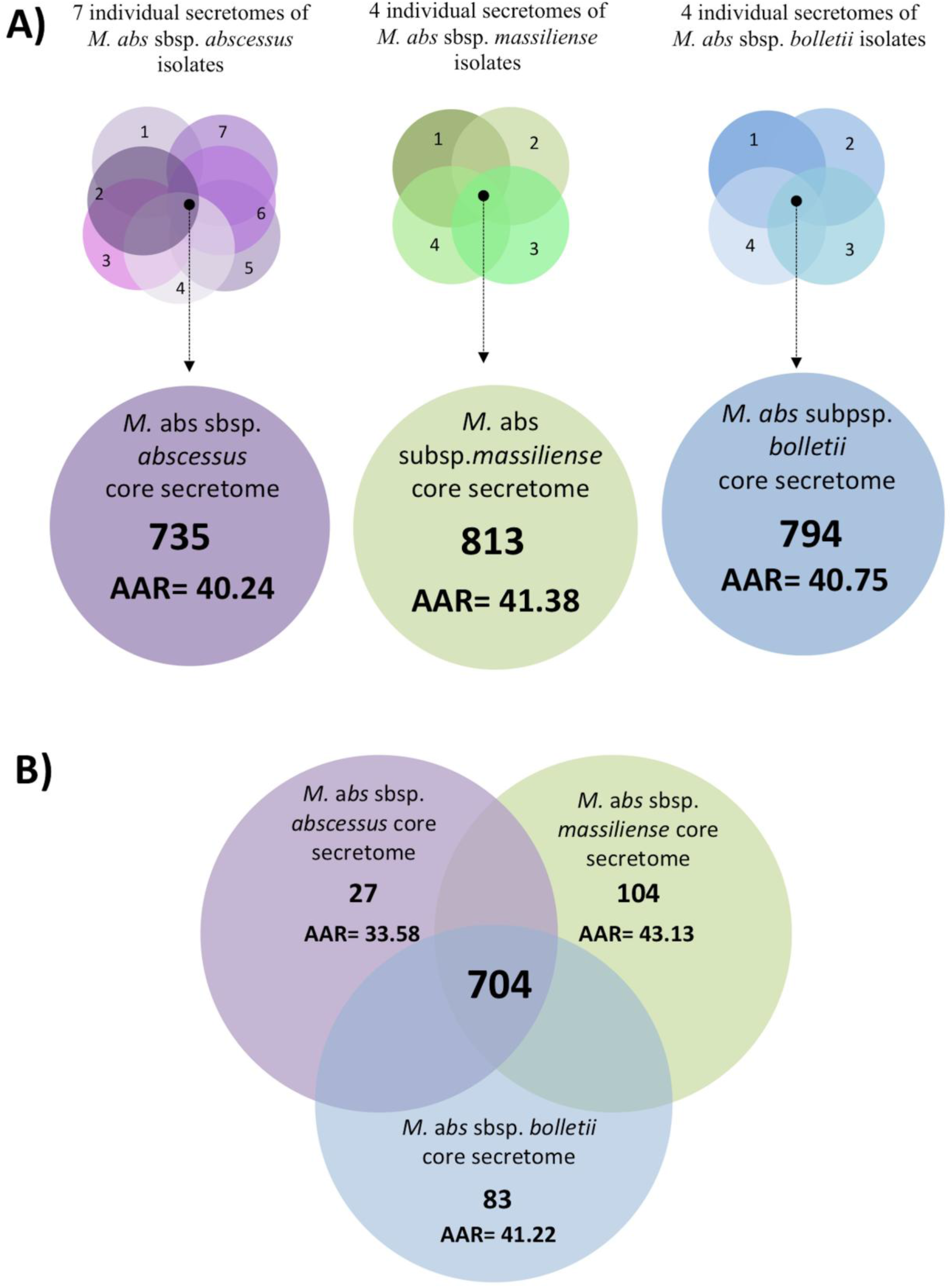
Venn diagram between the core secretomes of the three *M. abscessus subspecies*. A) Number of total proteins contained in the core secretome of each subspecies. B) Shared and unique proteins between the three subspecies as per BLASTP (E-value 1.0E-3).

Given that our study considered a limited number of de novo assembled genomes, we additionally compared the predicted core secretomes to sixty additional MAB genomes available in NCBI (Supplementary Table S3). We found that an average of 99.78%, 99.12%, and 98.59% of our core secretomes was also present in the investigated additional MABA, MABB, and MABM genomes, respectively, further corroborating the validity of the predicted subspecies core secretomes for other MAB isolates.

We then determined the respective AAR values to estimate antigenic densities for the protein sets in each core secretome. The average AAR values from most to least antigenic was: 40.24 for MABA, 40.75 for MAB_B_, and 41.38 for MAB_M_ with no statistically significant difference between these values.

Next, we identified the ES proteins shared between the MABA, MABB, and MABM core secretomes. We found that 704 proteins (86.5 %) were shared among MABA, MABB, and MAB_M_ with an AAR value of 41.17 (Fig. 2B). The AAR values for the protein sets exclusively found in the MAB_A_, MAB_B_, or MAB_M_ secretome were 33.58, 41.22, and 43.13, respectively, with the MABA dataset showing a significantly lower AAR value indicating higher antigenicity than the others (p < 0.1; Fig. 2B).

### Differences in core secretomes between R and S morphotypes

As MAB isolates with R and S morphotypes show differences in virulence and pathogenicity, we compared the predicted core secretomes of R and S isolates (Fig. 3). We observed that the core secretomes of R variants were larger (840, 924 and 845 proteins for MABA, MABM, and MABB) than those of the investigated S variants (764, 872 and 833 proteins, respectively) with no significant differences in antigenic densities as per mean AAR values (Fig. 3). Intra-subspecies comparison of S and R secretomes revealed that 96.4 %, 90.7% and 95% of the identified ES proteins were found in both R and S morphotypes for MABA, MABM and MABB respectively. The number of unique proteins was larger in the core secretome of the R morphotypes (n = 93, 109, and 48 for MABA, MAB_M_, and MAB_B_) as compared to the S morphotypes (n = 9, 76, and 35, respectively; Fig.3). Interestingly, antigenic densities for the unique ES proteins of the R morphotypes were higher (AAR = 40.84, 36.71, and 35.59 for MABA, MAB_M_, and MAB_B_) than for the proteins exclusive to the S morphotypes irrespective of the subspecies (AAR = 45.43, 37.72, and 42.14; Fig. 3). To assess if the AAR values of these specific protein sets were different from same-sized protein sets randomly chosen from the respective core secretomes, we created 1000 random sets of 109, 93, 76, 48, 35 and 9 proteins and calculated the AAR value for each set. Then, we determined an empirical p-value based on the number of random protein sets that equaled or exceeded the AAR value for each protein dataset as was previously suggested by Cornejo-Granados et al., 2017. We found that the ES proteins exclusive to the R morphotypes of MABM and MABB had significantly (p <0.05) higher antigenic densities than randomly constructed protein sets (Supplementary Table S5).

**Figure 3.**
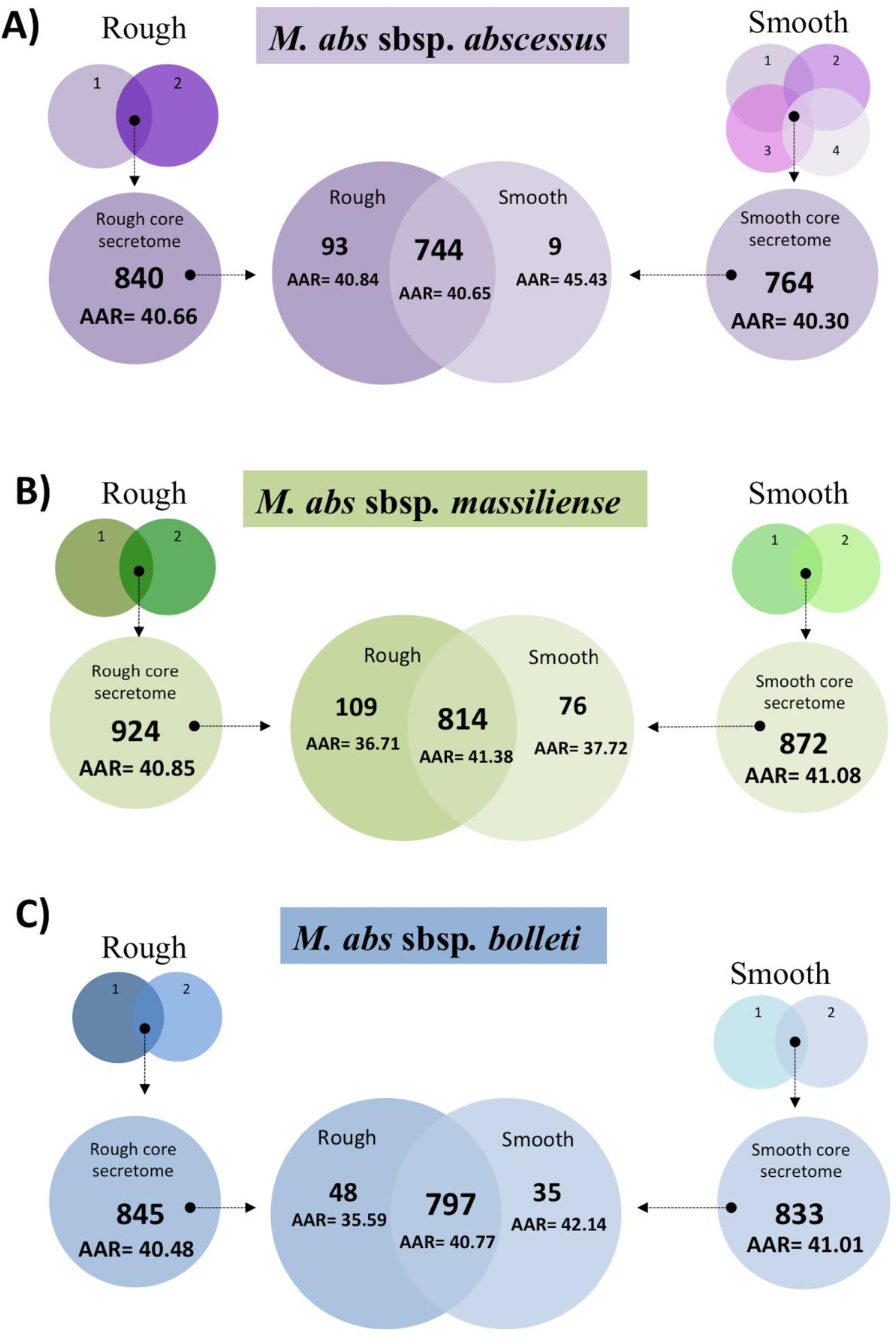
Venn diagram between the core secretomes of the three *M. abscessus* subspecies by colony morphotype. We used BLASTP (E-value 1.0E-3) to assess the core secretomes for isolates with rough and smooth colony morphotypes A) *M. abscessus* subsp. *abscessus*, B) *M. abscessus* subsp. *massiliense* and C) *M. abscessus* subsp. *bolletii*.

Finally, we determined the MAB core secretomes by sample origin (pulmonary, extrapulmonary, blood). This resulted in 706 ES proteins shared among the ten pulmonary isolates, 758 proteins shared among the four extrapulmonary isolates, and 885 proteins for the single isolate grown from a blood sample. However, as per the GO, KEGG, and antigenicity analyses, we did not find any distinct characteristics specific to either sample source and, hence, type of infection.

### Antigenicity of ES and non-ES proteins

It has previously been reported for different microorganisms including MTB that ES proteins tend to be more antigenic than non-ES proteins (Cornejo-Granados et al., 2017; Gomez et al., 2015; Wang et al., 2015). We thus tested if this was also true for the investigated MAB isolates. First, we found that the antigenic densities as indicated by mean AAR values were very similar among all isolates irrespective of subspecies or morphotype within the same cell compartment, i.e. for ES, non-ES, intracellular (incell) and transmembrane (TM) proteins (Fig. 4). Second, we found that antigenic densities were significantly higher in ES proteins as compared to non-ES proteins in all isolates (AAR = 40.57 and 43.60, respectively; p-value < 0.0001) (Fig. 4). However, within the non-ES category, incell proteins showed even higher antigenic densities (AAR = 39.04) than the predicted ES proteins (p < 0.0001) while the lowest overall antigenic densities were observed for the TM category (AAR = 59.23; p < 0.0001).

**Figure 4.**
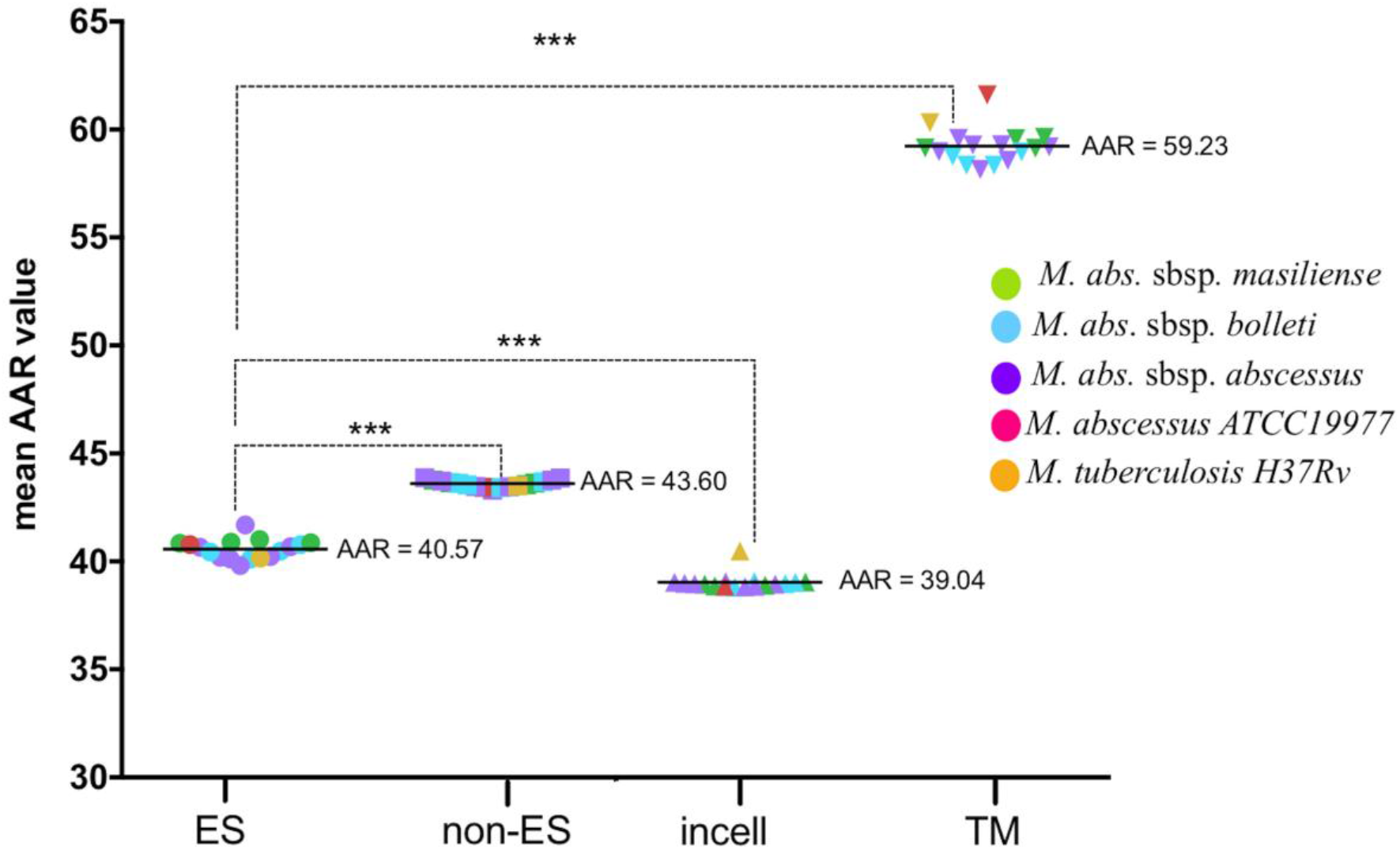
Comparison between AAR values for Excreted/Secreted (ES), non Excreted/Secreted (non-ES), intracellular (incell) and transmembrane (TM) proteins. AAR values were calculated for each of the 15 genomes sequenced. The X-axis shows the cellular compartment and the Y-axis shows AAR values for the genomes of each subspecies: *M. abscessus* subsp. *abscessus* (green), *M. abscessus* subsp. *bolletii* (blue), *M. abscessus* subsp. *massiliense* (purple), *M. abscessus* ATCC19977 (red) and *M. tuberculosis* H37Rv (orange). Mann-Whitney tests were performed to compare the AAR of each group with a confidence level of 99% (***, p < 0.001).

### Comparison of *M. abscessus* and *M. tuberculosis* secretomes

Lastly, we compared the predicted secretome of *M. abscessus* ATCC 19977 against the previously reported secretome of *M. tuberculosis* H37Rv (Cornejo-Granados et al., 2017). We observed that the *M. abscessus* secretome was predicted to be almost equally antigenic (AAR=40.78) than the *M. tuberculosis* secretome (AAR=40.63) (Fig. 5). We found 337 MAB ES proteins (38.04%) with homology to proteins in the predicted MTB secretome (Fig. 5). In addition, 222 of these proteins had sequence homology with proteins experimentally reported as secreted in MTB (comparable experimental secretome data for MAB was not available to us) (Cornejo-Granados et al., 2017) (Supplementary Table S6). Furthermore, we determined the average AAR value of the 680 ES proteins shared among the fifteen MAB isolates (AAR = 41.53). This value means that antigenic density was lower than for the predicted secretome of *M. tuberculosis* H37Rv (AAR = 40.63) and two clinical *M. tuberculosis* isolates belonging to the Beijing lineage (isolate C3 AAR = 37.51 and isolate C4 AAR = 37.54,) (Table 4) (Cornejo-Granados et al., 2017). Finally, we identified 17 ES proteins with homologues in both MAB and *M. tuberculosis*, which are listed as targets for various FDA approved drugs (Supplementary Table S2).

**Figure 5.**
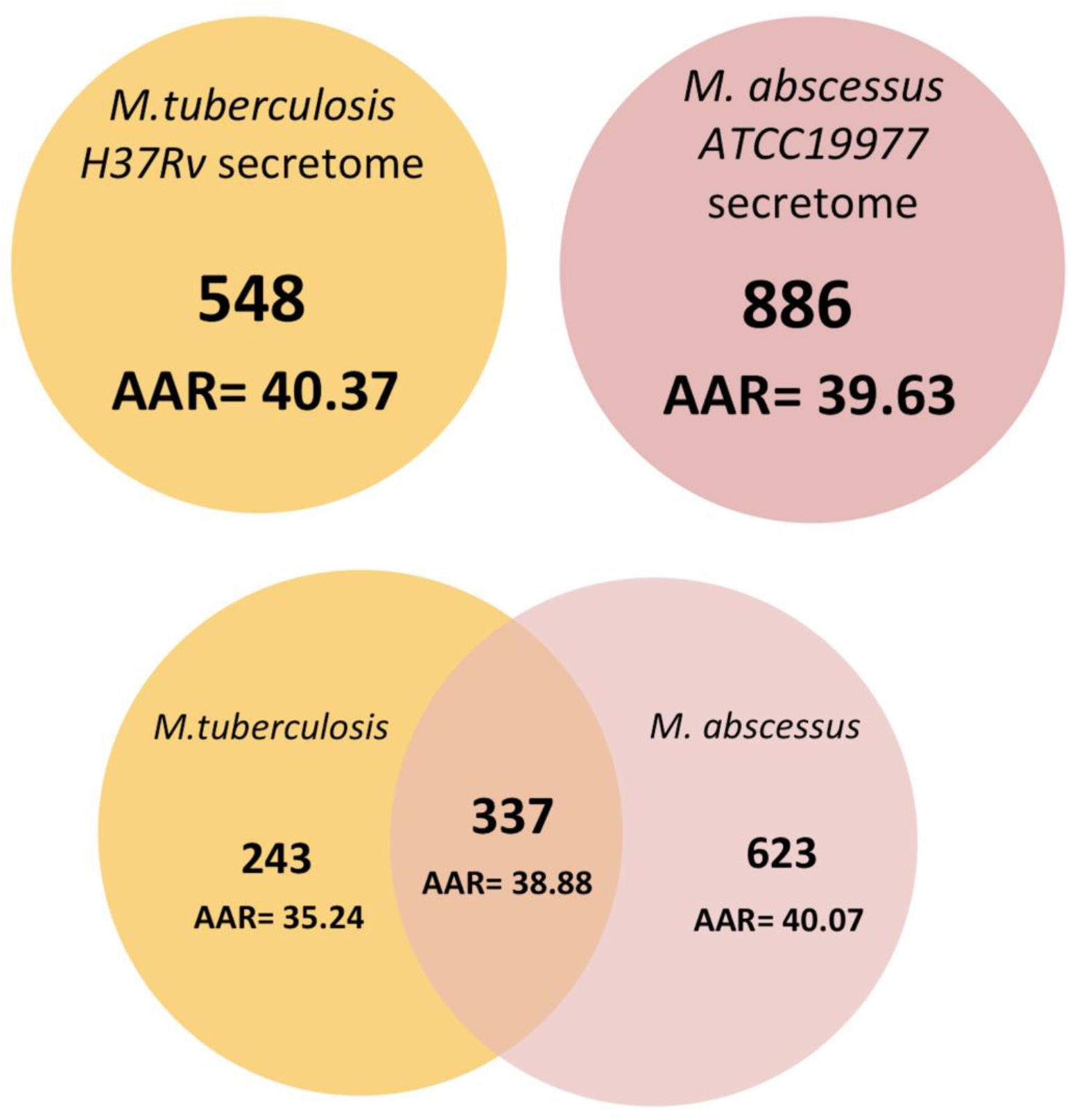
Venn diagram between the predicted secretomes of *M. tuberculosis* H37Rv and *M. abscessus* ATCC 19977. We used BLASTP (E-value 1.0E-3) to compare the complete secretomes of both species.

**Table 4.** Abundance of Antigenic Regions (AAR) for *M. abscessus* and *M. tuberculosis* strains.

| Strain | Number of proteins in the dataset | Average AAR value |
| --- | --- | --- |
| *M. tuberculosis* Beijing isolate C3* | 553 | 37.52 |
| *M. tuberculosis* Beijing isolate C4* | 519 | 37.55 |
| *M. bovis* BCG Pasteur | 564 | 38.99 |
| ***M. abscessus* ATCC 19977** | **886** | **40.78** |
| *M. tuberculosis* H37Rv | 548 | 40.63 |
| *M. abscessus* clinical isolates | 680 | 41.54 |

\* both Beijing isolates were previously reported in Cornejo-Granados et al., 2017.

## Discussion

This is the first study that proposes a method for prediction of MAB secretomes based on fifteen clinical MAB isolates and the *M. abscessus* ATCC 19977 reference strain. Our results show that an average of 18% (939 proteins) of the total proteins encoded in the MAB core genome carry sequence patterns indicative of secretion. Notably, this percentage is 6% larger than the proportion previously reported for several MTB isolates (~12%) (Cornejo-Granados et al., 2017). Nearly 200 species of mycobacteria have been identified with diverse lifestyles and a high degree of morphological, biochemical, and physiological diversity and a comparative genome analysis suggests that only a relatively small number of genes (1080) are shared between several *Mycobacterium* species (Malhotra et al., 2017; Tortoli et al., 2017). Moreover, loss of ancestral genes is a well described phenomenon in slowly growing mycobacteria such as MTB and, in particular, *M. leprae* (Bachmann et al., 2019). In contrast, rapidly growing NTM such as MAB are considered to represent a more ancient evolutionary state, with larger genomes than those of MTB (Bachmann et al., 2019; Malhotra et al., 2017). Thus, it is not surprising that we found a larger number of ES proteins in MAB than MTB. Furthermore, the increased abundance of ES proteins in MAB as compared to MTB could be related to the ability of MAB to cause a different spectrum of disease and to adapt to different environmental settings requiring frequent interaction with a wide variety of host cells and organisms competing for the same ecological niche, likely involving cross species exchange of genetic information, for example by plasmid transfer (Ripoll et al., 2009; Ryan and Byrd, 2018; Waman et al., 2019). A similar hypothesis has been suggested for fungal secretomes (O’Toole et al., 2013).

The GO and KEGG pathway annotations of the secretomes of *M. abscessus* ATCC 19977 and the MAB clinical isolates showed enrichment consistent with the characterization of previously reported mycobacterial secretomes (Cornejo-Granados et al., 2017; Gomez et al., 2015). Interestingly and in line with the increased secretome size as compared to MTB, the KEGG pathway analysis showed a high abundance of the Quorum sensing pathway for the predicted MAB secretomes, which was not present in our previous MTB secretome pathway analysis (Cornejo-Granados et al., 2017). The presence of a Quorum sensing pathway would be another similarity shared between MAB and non-mycobacterial pathogens commonly affecting patients with chronic lung disease such as *Pseudomonas aeruginosa* (Mukherjee and Bassler, 2019). In addition, it could be related to the ability of MAB to form biofilms (Clary et al., 2018; Orme and Ordway, 2014), further contributing to the capacity of MAB to tolerate antibiotics and to persist over long periods in the environment (Faria et al., 2015; Hunt-Serracin et al., 2019; Kulka et al., 2012; Maurer et al., 2014a).

The InterPro annotation showed that Mce domains were the most abundant (2.14 %) domains in the MAB reference secretome, while PPE and PE-PGRS domains only corresponded to 0.3 % of the ES protein sequences. This tendency is contrary to our observations for MTB (Cornejo-Granados et al., 2017), where the PPE and PE-PGRS domains accounted for ~12% of the secreted proteins and the Mce domains for only 0.5%. The lower quantity of predicted PE/PPE proteins in MAB was somewhat expected. *M. tuberculosis* has five ESX secretion systems, four of which encode PE/PPE proteins, while MAB has only two (ESX-3 and ESX-4) of which only the ESX-3 operon includes PE/PPE genes (Dumas et al., 2016). In contrast, Mce domains are known for participating in host cell entry by mycobacteria (Kumar et al., 2005). Thus, their higher abundance in MAB as compared to MTB highlights the importance of this pathway for MAB survival within the host. It needs to be mentioned though that Kumar et al., 2005, also suggested that in low virulence bacteria, transport activities could be the primary function of Mce operons (Kumar et al., 2005).

To compare the predicted secretomes according to colony morphotype, we first established the core secretome for the R and S variants per subspecies, thus eliminating individualities among the different isolates (Fig. 3). The high overall agreement between the core secretomes for both morphotypes of approximately 90 % was expected, considering the fact that R variants can arise from the S morphotypes during persistent infection by loss of surface-exposed GPLs caused by mutations in the GPL synthesis pathway (Bernut et al., 2014; Catherinot et al., 2007; Roux et al., 2016; Ryan and Byrd, 2018). However, both the higher number and the higher antigenic densities (lower AAR values) of the ES proteins exclusively found in R variants indicate that additional genetic changes may evolve during S-to-R conversion. Moreover, this observation raises the question whether some strains with additional genetic traits associated with virulence are able to undergo S-to-R conversion and cause disease due to R variants more easily than others. Genomic studies involving sequentially isolated S and R variants of the same strain obtained from individual patients over time will be required to better characterize the microevolution of MAB strains within the chronically infected host.

Similarly, the fact that MAB causes both chronic pulmonary disease (with R variants sometimes increasing over time) and extrapulmonary manifestations (mostly caused by S variants) led us to investigate whether differences exist in the predicted secretomes of isolates related to these clinical presentations. The absence of major differences in the GO, KEGG, and antigenicity analyses suggest that secretome variations do not influence MAB tissue tropism. Consequently, host characteristics such as severe immunosuppression may be the main driver for invasive MAB infections. Likewise, in the case of tissue infections, which often occur following surgical interventions, insufficient hygiene procedures and sterilization protocols for surgical equipment appear to be more relevant than pathobiological traits such as the secretome intrinsic to the causative MAB isolate (Maurer et al., 2014b)

Lastly, we observed that the predicted secretomes of all investigated clinical MAB isolates were less antigenic than the secretomes of *M. tuberculosis* H37Rv and two clinical *M. tuberculosis* isolates. Additionally, although there was no statistical difference, the isolates with a rough phenotype tended to be more antigenic that the isolates with smooth phenotype. Previous evidence with *M. tuberculosis* (Cornejo-Granados et al., 2017) showed that clinical isolates from the Beijing phenotype showed increased virulence and less antigenic secretomes than the reference strain H37Rv. Thus, the diminished antigenicity of MAB could be viewed as a virulence trait in itself as it would support colonization of the host for extended time periods without immediate progression into clinical disease. However, further experimental tests on antigenicity are needed to demonstrate this observation.

This study represents the first systematic prediction and *in silico* characterization of the MAB secretome. We acknowledge that an important constraint of this study is the limited total number of genomes analyzed per subspecies and biological source. Thus, care must be taken to not overinterpret the findings related to sample subcategories such as subspecies and morphotypes. Also, published experimental data on MAB secretomes are very limited and no systematic validation of the *in silico* findings reported herein could be performed against such datasets. Although, more research will be needed to determine experimental secretomes in NTM, our study demonstrates that using bioinformatics strategies can help to broadly explore mycobacterial secretomes including those of clinical isolates and to tailor subsequent, complex and time-consuming experimental approaches accordingly. This approach can support a systematic investigation of mycobacterial secretomes exploring candidate proteins suitable for developing new vaccines and diagnostic markers to distinguish between colonization and infection.

## Supporting information

Data Sheet 1

Data Sheet 2

## Acknowledgements

We would like to thank Julia Zallet and Vanessa Mohr for excellent technical assistance and Henrik Nielsen for his assistance with the SecrtomeP software. Parts of this work have been supported by the German Center for Infection Research and Grants by Joachim Herz Foundation, Hamburg, and Mukoviszidose Institut gGmbH, Bonn, the research and development arm of the German Cystic Fibrosis Association Mukoviszidose e.V to F.P.M. We acknowledge the support provided by CONACyT grant CB-2013-223279 and SALUD-2014-C01-234188 to A.O.L. This research also received support by the DGAPA PAPIIT UNAM (IN215520) to A.O.L. F.C.G. acknowledges the support of CONACyT as a Postgraduate fellow.

