## Supplementary material for "*In silico* secretome characterization of clinical *Mycobacterium abscessus* isolates provides insights into antigenic differences": Data Sheet 1

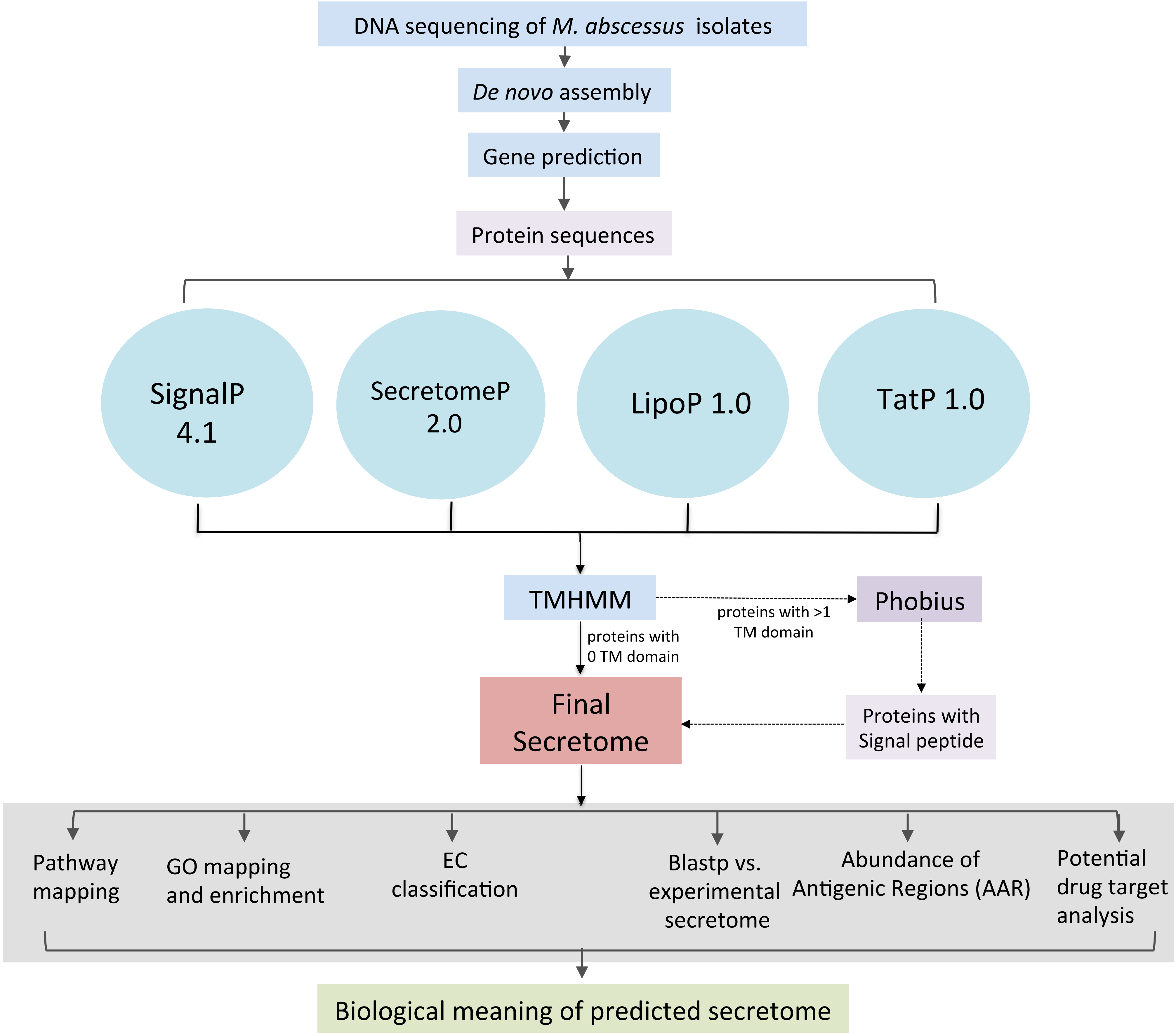


**Supplementary Figure S1**. Bioinformatics pipeline to identify and analyze the secreted proteins of *M. abscessus*.

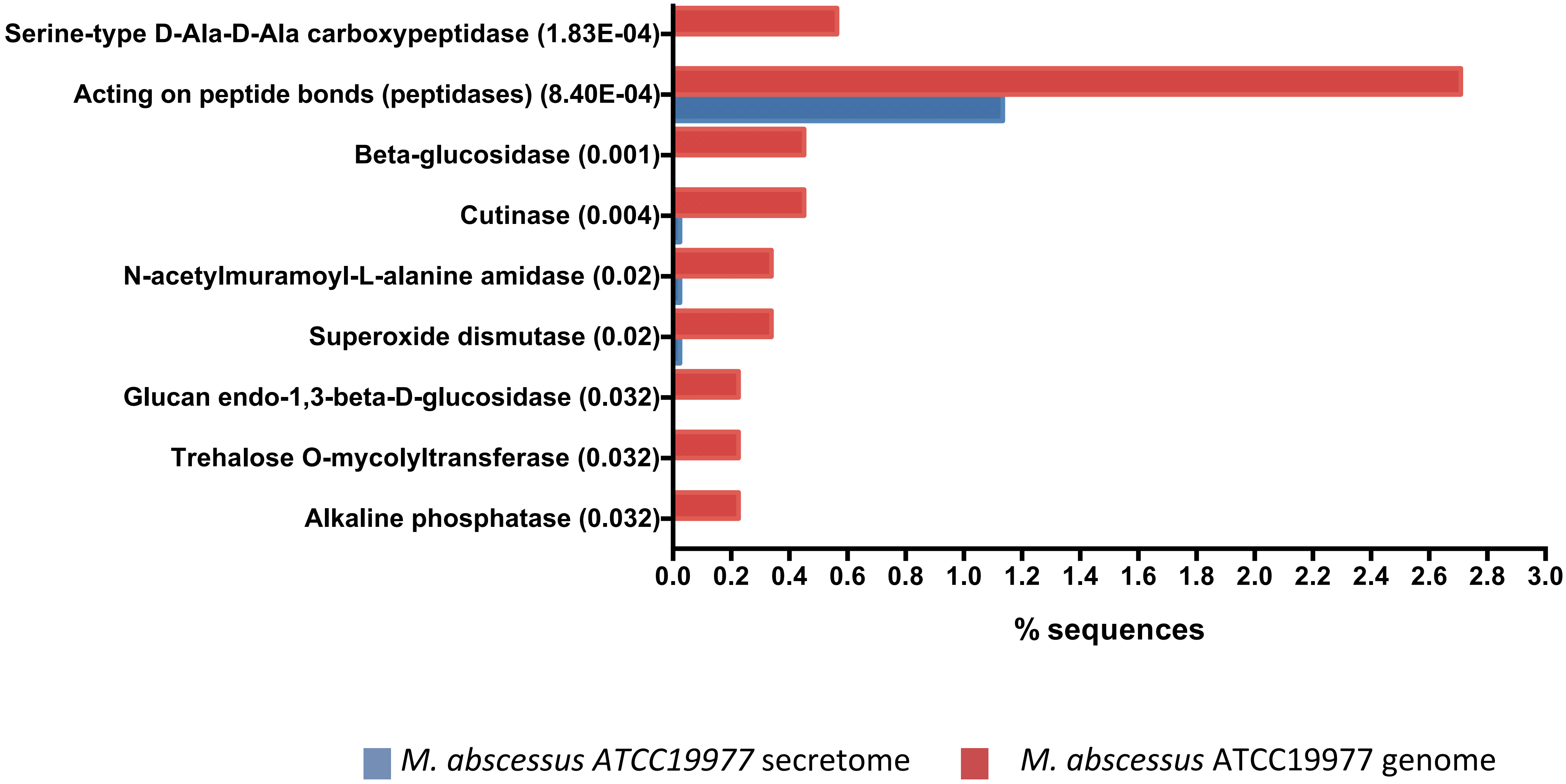


**Supplementary Fig S2. GO enrichment analysis of Enzymes for the *M. abscessus* ATCC19977.** Percentage of sequences annotated with each GO term for the secretome proteins (blue) and the complete proteins in the genome (red).
