## Supplementary material for "*In silico* secretome characterization of clinical *Mycobacterium abscessus* isolates provides insights into antigenic differences": Data Sheet 2

Table S1. Complete metadata of the fifteen clinical isolates of *M. abscessus* genomes sequenced.

| **Species** | **Genome ID** | **Origin** | **Phenotype** | **Total predicted proteins** | **ES proteins** | **non-ES proteins** | **incell proteins** | **TM proteins** | **mean ES AAR** | **mean non-ES AAR** | **mean incell AAR** | **mean TM AAR** |
|---|---|---|---|---|---|---|---|---|---|---|---|---|
| *M. abscessus sbsp. abscessus* | 4549-15 | sputum | rough | 5,105 | 929 | 4,176 | 3,213 | 963 | 40.66 | 43.48 | 39.07 | 58.34 |
| | 11351-15 | sputum | rough | 5,138 | 966 | 4,172 | 3,215 | 957 | 40.12 | 43.63 | 39.05 | 59.20 |
| | 8844-15 | skin | smooth | 4,854 | 956 | 3,898 | 2,974 | 924 | 39.81 | 43.87 | 39.04 | 59.58 |
| | 3563-15 | sputum | smooth | 5,239 | 968 | 4,271 | 3,271 | 1,000 | 40.11 | 43.43 | 38.99 | 58.15 |
| | 12389-15 | sputum | smooth | 5,276 | 990 | 4,286 | 3,314 | 972 | 40.19 | 43.30 | 38.95 | 58.35 |
| | 2677-16 | sputum | smooth | 4,900 | 919 | 3,981 | 3,024 | 957 | 40.68 | 43.87 | 39.06 | 59.28 |
| | 2572-17 | tissue (breast implant) | NA | 4,847 | 874 | 3,973 | 3,039 | 934 | 40.47 | 43.76 | 38.98 | 59.58 |
| *M. abscessus sbsp. massiliense* | 14479-15 | sputum | rough | 5,120 | 962 | 4,158 | 3,190 | 968 | 40.89 | 43.65 | 38.85 | 59.64 |
| | 10896-16 | sputum | rough | 5,109 | 950 | 4,159 | 3,192 | 967 | 40.86 | 43.54 | 38.86 | 59.15 |
| | 10003-15 | sputum | smooth | 4,835 | 891 | 3,944 | 3,017 | 927 | 41.01 | 43.61 | 38.86 | 59.30 |
| | 16155-15 | sputum | smooth | 4,884 | 898 | 3,986 | 3,061 | 925 | 40.84 | 43.50 | 38.83 | 59.14 |
| *M. abscessus sbsp. bolletii* | 11702-16 | sputum | rough | 5,079 | 931 | 4,148 | 3,177 | 971 | 40.22 | 43.56 | 38.90 | 58.97 |
| | 713-16 | lymph node | rough | 5,456 | 1,037 | 4,419 | 3,401 | 1,018 | 40.78 | 43.42 | 38.95 | 58.57 |
| | 7742-15 | blood culture | smooth | 4,913 | 885 | 4,028 | 3,067 | 961 | 41.69 | 43.69 | 39.04 | 58.77 |
| | 13116-16 | lymph node | smooth | 5,305 | 990 | 4,315 | 3,312 | 1,003 | 40.44 | 43.46 | 38.82 | 58.96 |
| *M. abscessus* | reference strain ATCC19977 (GenBank CP016193.1) | - | - | 4,942 | 886 | 4,056 | 3,196 | 860 | 40.78 | 43.72 | 38.96 | 61.60 |

Supplementary Table S2. Potential drug targets for the 222 proteins shared between *M. tuberculosis H37Rv* and *M. abscessus* ATCC19977.

| Protein | Target | E value | Approved drug |
|---|---|---|---|
| YP_001700862.1 | Protein disulfide-isomerase A3 | 3.33E-09 | Copper, Zinc, Zinc acetate, Zinc chloride |
| YP_001700866.1 | D-beta-hydroxybutyrate dehydrogenase, mitochondrial | 4.59E-16 | NADH |
| YP_001700902.1 | Retinol dehydrogenase 8 | 3.28E-15 | Vitamin A |
| YP_001700905.1 | Estradiol 17-beta-dehydrogenase 1 | 1.98E-14 | NADH, Prasterone |
| YP_001700922.1 | Estradiol 17-beta-dehydrogenase 2 | 7.36E-12 | NADH, Estradiol, Estradiol acetate, Estradiol benzoate, Estradiol cypionate, Estradiol dienanthate, Estradiol valerate |
| YP_001700923.1 | Dehydrogenase/reductase SDR family member 4 | 5.55E-10 | Vitamin A |
| YP_001701031.1 | Short-chain dehydrogenase/reductase 3 | 9.07E-10 | Vitamin A |
| YP_001701057.1 | Corticosteroid 11-beta-dehydrogenase isozyme 1 | 4.97E-09 | NADH |
| YP_001701074.1 | Estradiol 17-beta-dehydrogenase 8 | 8.97E-08 | NADH |
| YP_001701078.1 | 17-beta-hydroxysteroid dehydrogenase type 6 | 1.30E-07 | Succinic acid |
| YP_001701111.1 | 15-hydroxyprostaglandin dehydrogenase [NAD(+)] | 1.44E-07 | NADH |
| YP_001701113.1 | Peroxisomal multifunctional enzyme type 2 | 4.06E-07 | NADH |
| YP_001701114.1 | Corticosteroid 11-beta-dehydrogenase isozyme 2 | 1.64E-06 | NADH, Fluoxymesterone, Hydrocortisone, Hydrocortisone acetate, Hydrocortisone butyrate, Hydrocortisone cypionate, Hydrocortisone phosphate, Hydrocortisone probutate |
| YP_001701121.1 | 11-cis retinol dehydrogenase | 8.68E-06 | NADH, Vitamin A |
| YP_001701129.1 | 3-hydroxyacyl-CoA dehydrogenase type-2 | 3.37E-05 | NADH, Omega-3-carboxylic acids |
| YP_001701177.1 | Superoxide dismutase [Cu-Zn] | 3.55E-07 | Vitamin E, Cisplatin, Carboplatin, Oxaliplatin, Copper, Zinc, Dopamine |
| YP_001701206.1 | Copper chaperone for superoxide dismutase | 5.00E-06 | Copper, Zinc, Zinc acetate, Zinc chloride |
| YP_001701209.1 | N-acetylmuramoyl-L-alanine amidase | 1.10E-11 | Copper, Zinc, Zinc acetate, Zinc chloride |
| YP_001701213.1 | Thiosulfate sulfurtransferase | 2.32E-27 | Thiosulfuric acid |
| YP_001701267.1 | Deazaflavin-dependent nitroreductase | 2.10E-52 | Delamanid, Pretomanid |
| YP_001701292.1 | Retinol dehydrogenase 14 | 2.68E-12 | Vitamin A |
| YP_001701361.1 | Retinol dehydrogenase 12 | 6.78E-08 | Vitamin A |
| YP_001701380.1 | Retinol dehydrogenase 13 | 6.88E-08 | Vitamin A |
| YP_001701382.1 | Carbapenem-hydrolyzing beta-lactamase KPC | 6.84E-81 | Avibactam, Vaborbactam |
| YP_001701410.1 | Beta-lactamase | 4.16E-80 | Citric acid, Cefoxitin |
| YP_001701416.1 | Beta-lactamase UOE-1 | 4.27E-73 | Avibactam |
| YP_001701434.1 | Beta-lactamase SHV-1 | 1.91E-54 | Avibactam, Tazobactam, Relebactam, Imipenem |
| YP_001701435.1 | Peptidyl-prolyl cis-trans isomerase B | 7.48286e-28 | Proline |
| YP_001701467.1 | Peptidyl-prolyl cis-trans isomerase A | 5.68E-22 | Cyclosporine |
| YP_001701472.1 | Deazaflavin-dependent nitroreductase | 5.09E-21 | Delamanid, Pretomanid |
| YP_001701473.1 | Bile salt export pump | 2.91E-13 | Glyburide, Ursodeoxycholic acid, Deoxycholic acid, Taurocholic acid, Cholic Acid, Dexamethasone, Phenobarbital, Ethinylestradiol, Rifampicin |
| YP_001701484.1 | Phosphatidylcholine translocator ABCB4 | 1.73822e-12 | Silodosin, Etravirine |
| YP_001701485.1 | ATP-binding cassette sub-family B member 8, mitochondrial | 1.06E-10 | Doxorubicin |
| YP_001701486.1 | ATP-binding cassette sub-family B member 5 | 3.95E-10 | Dasabuvir, Delafloxacin, Bilastine, Naldemedine, Vinflunine, Brentuximab, Fostamatinib |
| YP_001701490.1 | ATP-binding cassette sub-family A member 3 | 1.14E-06 | Imatinib |
| YP_001701496.1 | Antigen peptide transporter 1 | 1.38E-06 | Lapatinib |
| YP_001701525.1 | ATP-binding cassette sub-family A member 1 | 2.46E-06 | Glyburide, Probucol, Tocofersolan, Vitamin E, Tamoxifen |
| YP_001701528.1 | ATP-binding cassette sub-family C member 11 | 3.35E-06 | Indomethacin, Probenecid, Taurocholic acid, Conjugated estrogens, Folic acid, Methotrexate |
| YP_001701579.1 | Canalicular multispecific organic anion transporter 1 | 4.36E-06 | Sulfinpyrazone, Ritonavir, Ursodeoxycholic acid, Rifampicin, Arsenic trioxide, Cisplatin |
| YP_001701589.1 | Canalicular multispecific organic anion transporter 2 | 1.48E-05 | Nifedipine, Metyrapone, Sulfinpyrazone, Probenecid, Indomethacin, Vincristine |
| YP_001701617.1 | ATP-binding cassette sub-family C member 9 | 3.85E-05 | Glyburide, Nicorandil |
| YP_001701632.1 | 3-hydroxyacyl-CoA dehydrogenase type-2 | 6.97E-68 | Omega-3-carboxylic acids |
| YP_001701633.1 | Peroxisomal trans-2-enoyl-CoA reductase | 6.15E-08 | Adenine |
| YP_001701713.1 | 17-beta-hydroxysteroid dehydrogenase type 6 | 5.62E-12 | Succinic acid |
| YP_001701720.1 | Estradiol 17-beta-dehydrogenase 1 | 2.42E-10 | Prasterone |
| YP_001701724.1 | RNA polymerase sigma factor SigA | 1.05E-113 | Fidaxomicin |

Supplementary Table S3 | Comparisson of the core secretome of each subspecies vs NCBI genomes.

| | GenBank Accession number | Numer of ES core proteins present in the new genomes | % of ES core proteins present in the new genomes | Average % |
|---|---|---|---|---|
| *Comparisson to M. abscessus* sbsp. *abscessus genomes (735 core ES proteins)* | GCA_900141265.1 | 734 | 99.86 | 99.78 |
| | GCA_900140885.1 | 733 | 99.73 | |
| | GCA_900140835.1 | 733 | 99.73 | |
| | GCA_900140825.1 | 732 | 99.59 | |
| | GCA_900140545.1 | 732 | 99.59 | |
| | GCA_900140475.1 | 733 | 99.73 | |
| | GCA_900140045.1 | 734 | 99.86 | |
| | GCA_900140005.1 | 734 | 99.86 | |
| | GCA_900139935.1 | 732 | 99.59 | |
| | GCA_900139505.1 | 734 | 99.86 | |
| | GCA_900137175.1 | 733 | 99.73 | |
| | GCA_900137165.1 | 734 | 99.86 | |
| | GCA_900137155.1 | 734 | 99.86 | |
| | GCA_900137145.1 | 734 | 99.86 | |
| | GCA_900136765.1 | 734 | 99.86 | |
| | GCA_900136595.1 | 734 | 99.86 | |
| | GCA_900135095.1 | 733 | 99.73 | |
| | GCA_900135045.1 | 734 | 99.86 | |
| | GCA_900132315.1 | 733 | 99.73 | |
| | GCA_900132295.1 | 734 | 99.86 | |
| Comparisson to M. abscessus sbsp. bolletii genomes (794 core ES proteins) | GCA_900132985.1 | 778 | 97.98 | 99.12 |
| | GCA_900133045.1 | 790 | 99.50 | |
| | GCA_900133055.1 | 788 | 99.24 | |
| | GCA_900133105.1 | 788 | 99.24 | |
| | GCA_900133625.1 | 788 | 99.24 | |
| | GCA_900133635.1 | 788 | 99.24 | |
| | GCA_900133785.1 | 787 | 99.12 | |
| | GCA_900134275.1 | 785 | 98.87 | |
| | GCA_900134285.1 | 784 | 98.74 | |
| | GCA_900134335.1 | 787 | 99.12 | |
| | GCA_900134535.1 | 787 | 99.12 | |
| | GCA_900135035.1 | 790 | 99.50 | |
| | GCA_900136205.1 | 792 | 99.75 | |
| | GCA_900136235.1 | 792 | 99.75 | |
| | GCA_900136655.1 | 789 | 99.37 | |
| | GCA_900137535.1 | 789 | 99.37 | |
| | GCA_900137885.1 | 786 | 98.99 | |
| | GCA_900139995.1 | 781 | 98.36 | |
| | GCA_900141605.1 | 784 | 98.74 | |
| | GCF_900131565.1 | 787 | 99.12 | |
| Comparisson to M. abscessus sbsp. massiliense genomes (813 core ES proteins) | GCA_900130325.1 | 805 | 99.02 | 98.59 |
| | GCA_900130675.1 | 802 | 98.65 | |
| | GCA_900130855.1 | 804 | 98.89 | |
| | GCA_900131585.1 | 798 | 98.15 | |
| | GCA_900134075.1 | 803 | 98.77 | |
| | GCA_900135905.1 | 804 | 98.89 | |
| | GCA_900135915.1 | 802 | 98.65 | |
| | GCA_900137185.1 | 803 | 98.77 | |
| | GCA_900138505.1 | 802 | 98.65 | |
| | GCA_900138535.1 | 802 | 98.65 | |
| | GCA_900138705.1 | 806 | 99.14 | |
| | GCA_900138715.1 | 806 | 99.14 | |
| | GCA_900139775.1 | 801 | 98.52 | |
| | GCA_900139915.1 | 797 | 98.03 | |
| | GCA_900140635.1 | 795 | 97.79 | |
| | GCA_900140755.1 | 795 | 97.79 | |
| | GCA_900141015.1 | 800 | 98.40 | |
| | GCA_900141505.1 | 801 | 98.52 | |
| | GCA_900141515.1 | 803 | 98.77 | |
| | GCA_900169205.1 | 802 | 98.65 | |

Table S4.| Statistic data of the de novo assemblies for the sequenced isolates.

| **Genome** | **Strain** | **Total reads** | **N50** | **Mean base coverage** | **Largest Contig** | **Total Contigs** |
|---|---|---|---|---|---|---|
| 4549-15 | *M. abscessus* | 1,841,762 | 241,795 | 217.25 | 479,260 | 61 |
| 11351-15 | *M. abscessus* | 2,405,256 | 200,998 | 284.07 | 455,684 | 75 |
| 8844-15 | *M. abscessus* | 2,483,126 | 220,207 | 307.33 | 366,033 | 60 |
| 3563-15 | *M. abscessus* | 2,941,828 | 218,353 | 343.09 | 775,036 | 52 |
| 12389-15 | *M. abscessus* | 2,241,946 | 232,848 | 257.56 | 357,818 | 63 |
| 2677-16 | *M. abscessus* | 2,755,836 | 347,263 | 337.93 | 701,340 | 71 |
| 2572-17 | *M. abscessus* | 2,576,134 | 337,055 | 317.67 | 1,047,246 | 38 |
| 11702-16 | *M. bolletii* | 2,255,592 | 652,527 | 268.14 | 1,269,368 | 53 |
| 713-16 | *M. bolletii* | 2,641,622 | 384,872 | 296.01 | 607,861 | 68 |
| 7742-15 | *M. bolletii* | 2,989,318 | 302,394 | 365.44 | 962,231 | 47 |
| 13116-16 | *M. bolletii* | 2,785,738 | 344,725 | 319.87 | 1,196,314 | 55 |
| 14479-15 | *M. massiliense* | 3,100,556 | 327,410 | 368.37 | 722,347 | 64 |
| 10896-16 | *M. massiliense* | 2,510,338 | 327,557 | 298.78 | 522,805 | 47 |
| 10003-15 | *M. massiliense* | 2,667,154 | 511,575 | 329.83 | 1,346,120 | 39 |
| 16155-15 | *M. massiliense* | 2,825,456 | 339,641 | 345.85 | 1,087,576 | 78 |

Table S5.| AAR values for random constructed secretomes of the rough and smooth phenotypes.

| | Number of proteins in the set | empirical *p* value |
|---|---|---|
| *M. abs* sbsp. *massiliense* unique rough proteins | 109 | 0.01 |
| *M. abs* sbsp. *bolletii* unique rough proteins | 48 | 0.04 |
| *M. abs* sbsp. *abscessus* unique smooth proteins | 9 | 0.19 |
| *M. abs* sbsp. *bolletii* unique smooth proteins | 35 | 0.37 |
| *M. abs* sbsp. *abscessus* unique rough proteins | 93 | 0.58 |
| *M. abs* sbsp. *massiliense* unique smooth proteins | 76 | 0.93 |

sets with significative $p$ value <0.05

Supplementary Table S6 | List of 222 *M. abscessus* proteins with homologues in *M. tuberculosis* H37Rv and with previous experimental support for secretion according to [Cornejo-Granados et al., 2017 paper]".

| | Protein name | Description |
|---|---|---|
| 1 | YP_001700711.1 | Conserved hypothetical protein (plasmid) |
| 2 | YP_001700767.1 | Hypothetical protein MAB_0010c |
| 3 | YP_001700781.1 | Peptidyl-prolyl cis-trans isomerase |
| 4 | YP_001700795.1 | Conserved hypothetical protein |
| 5 | YP_001700847.1 | Conserved hypothetical protein |
| 6 | YP_001700862.1 | Hypothetical protein MAB_0108c |
| 7 | YP_001700866.1 | Hypothetical protein MAB_0112 |
| 8 | YP_001700902.1 | PPE family protein |
| 9 | YP_001700905.1 | Conserved hypothetical protein |
| 10 | YP_001700922.1 | Putative N-acetymuramoyl-L-alanine amidase |
| 11 | YP_001700923.1 | Putative exported repetitive protein precursor |
| 12 | YP_001701031.1 | Probable amino acid ABC transporter, permease |
| 13 | YP_001701057.1 | Hypothetical protein MAB_0304 |
| 14 | YP_001701074.1 | Hypothetical protein MAB_0321c |
| 15 | YP_001701078.1 | Hypothetical protein MAB_0325c |
| 16 | YP_001701111.1 | Hypothetical conserved membrane protein PknM or conserved lipoprotein LppH |
| 17 | YP_001701113.1 | Conserved hypothetical protein |
| 18 | YP_001701114.1 | Conserved hypothetical protein |
| 19 | YP_001701121.1 | Short-chain dehydrogenase/reductas |
| 20 | YP_001701129.1 | Conserved hypothetical protein (peptidase?) |
| 21 | YP_001701177.1 | Putative protease |
| 22 | YP_001701206.1 | Conserved hypothetical protein |
| 23 | YP_001701209.1 | Hypothetical protein MAB_0456 |
| 24 | YP_001701213.1 | Hypothetical protein MAB_0460 |
| 25 | YP_001701267.1 | Probable oxidoreductase EphD |
| 26 | YP_001701292.1 | Conserved hypothetical transmembrane protein |
| 27 | YP_001701361.1 | Conserved hypothetical protein |
| 28 | YP_001701380.1 | Conserved hypothetical protein |
| 29 | YP_001701382.1 | Conserved hypothetical protein |
| 30 | YP_001701410.1 | Conserved hypothetical protein |
| 31 | YP_001701416.1 | PE family protein |
| 32 | YP_001701434.1 | Hypothetical conserved membrane protein |
| 33 | YP_001701435.1 | Hypothetical conserved protein |
| 34 | YP_001701467.1 | Probable dehydrogenase/reductase |
| 35 | YP_001701472.1 | Putative oligopeptide ABC transporter, ATP-binding protein |
| 36 | YP_001701473.1 | Putative triacylglycerol lipase precursor |
| 37 | YP_001701484.1 | Conserved hypothetical protein |
| 38 | YP_001701485.1 | Hypothetical protein MAB_0735 |
| 39 | YP_001701486.1 | Hypothetical protein MAB_0736 |
| 40 | YP_001701490.1 | Probable thiosulfate sulfurtransferase (CysA) |
| 41 | YP_001701496.1 | Putative phosphate ABC transporter, phosphate-binding protein |
| 42 | YP_001701525.1 | Hypothetical protein MAB_0775 |
| 43 | YP_001701528.1 | Conserved hypothetical protein (lipoprotein LppU?) |
| 44 | YP_001701579.1 | Conserved hypothetical protein |
| 45 | YP_001701589.1 | Hypothetical protein MAB_0841 |
| 46 | YP_001701617.1 | Probable resuscitation-promoting factor RpfA |
| 47 | YP_001701632.1 | Hypothetical protein MAB_0884c |
| 48 | YP_001701633.1 | Hypothetical lipoprotein lpqH precursor |
| 49 | YP_001701713.1 | Hypothetical protein MAB_0967 |
| 50 | YP_001701720.1 | Conserved hypothetical protein |
| 51 | YP_001701724.1 | Hypothetical protein MAB_0978 |
| 52 | YP_001701750.1 | Putative MCE family protein |
| 53 | YP_001701753.1 | Putative MCE family protein |
| 54 | YP_001701754.1 | Putative MCE family protein |
| 55 | YP_001701757.1 | Hypothetical protein MAB_1013 |
| 56 | YP_001701770.1 | Conserved hypothetical protein |
| 57 | YP_001701857.1 | Hypothetical protein MAB_1115 |

| | Protein name | Description |
|---|---|---|
| 58 | YP_001701872.1 | Conserved hypothetical protein |
| 59 | YP_001701906.1 | Putative conserved lipoprotein LpqU |
| 60 | YP_001701920.1 | Conserved hypothetical protein |
| 61 | YP_001701935.1 | Conserved hypothetical protein (lipolytic enzyme G-D-S-L?) |
| 62 | YP_001701943.1 | Hypothetical transcription elongation factor GreA |
| 63 | YP_001702022.1 | Hypothetical protein MAB_1280c |
| 64 | YP_001702045.1 | Hypothetical protein MAB_1303c |
| 65 | YP_001702056.1 | Hypothetical protein MAB_1314 |
| 66 | YP_001702057.1 | Putative lipoprotein LpqW |
| 67 | YP_001702064.1 | Hypothetical protein MAB_1322 |
| 68 | YP_001702065.1 | Hypothetical protein MAB_1323 |
| 69 | YP_001702085.1 | Conserved hypothetical protein (nuclease?) |
| 70 | YP_001702107.1 | Conserved hypothetical protein |
| 71 | YP_001702142.1 | Putative lipoprotein LprE precursor |
| 72 | YP_001702156.1 | Putative lipoprotein LprB precursor |
| 73 | YP_001702157.1 | Putative lipoprotein LprC precursor |
| 74 | YP_001702177.1 | Probable homoserine kinase (ThrB) |
| 75 | YP_001702180.1 | Hypothetical protein MAB_1440c |
| 76 | YP_001702205.1 | Conserved hypothetical protein thioredoxin-like |
| 77 | YP_001702206.1 | Possible lipoprotein peptidase LpqM |
| 78 | YP_001702210.1 | Possible lipoprotein peptidase LpqM |
| 79 | YP_001702247.1 | Conserved hypothetical protein |
| 80 | YP_001702251.1 | Conserved hypothetical protein |
| 81 | YP_001702266.1 | Putative short chain dehydrogenase/reductase |
| 82 | YP_001702270.1 | Probable conserved lipoprotein LppS |
| 83 | YP_001702282.1 | Conserved hypothetical protein |
| 84 | YP_001702316.1 | Conserved hypothetical protein |
| 85 | YP_001702336.1 | Conserved hypothetical protein |
| 86 | YP_001702437.1 | Putative Mce family protein |
| 87 | YP_001702438.1 | Putative Mce family protein |
| 88 | YP_001702439.1 | Putative Mce family protein |
| 89 | YP_001702445.1 | Conserved hypothetical protein |
| 90 | YP_001702539.1 | Bacteriophage protein |
| 91 | YP_001702541.1 | Bacteriophage protein |
| 92 | YP_001702567.1 | Hypothetical protein MAB_1830 |
| 93 | YP_001702569.1 | Conserved hypothetical protein |
| 94 | YP_001702570.1 | Conserved hypothetical protein |
| 95 | YP_001702572.1 | Immunogenic protein MPT64 precursor |
| 96 | YP_001702577.1 | Putative beta-glucanase |
| 97 | YP_001702628.1 | Conserved hypothetical protein |
| 98 | YP_001702632.1 | Conserved hypothetical protein |
| 99 | YP_001702710.1 | Putative secreted protein |
| 100 | YP_001702717.1 | Conserved hypothetical protein |
| 101 | YP_001702838.1 | Conserved hypothetical protein |
| 102 | YP_001702895.1 | Putative lipoprotein LppK precursor |
| 103 | YP_001702896.1 | Hypothetical low molecular weight antigen Mtb12 |
| 104 | YP_001703063.1 | Conserved hypothetical protein |
| 105 | YP_001703114.1 | Hypothetical lipoprotein LpqH precursor |
| 106 | YP_001703138.1 | Conserved hypothetical protein |
| 107 | YP_001703155.1 | Conserved hypothetical protein |
| 108 | YP_001703156.1 | Conserved hypothetical protein |
| 109 | YP_001703168.1 | Molybdenum ABC transporter ModA, periplasmic |
| 110 | YP_001703171.1 | Conserved hypothetical protein (fibronectin-attachment?) |
| 111 | YP_001703191.1 | Conserved hypothetical protein |
| 112 | YP_001703209.1 | Conserved hypothetical protein |
| 113 | YP_001703223.1 | Conserved hypothetical protein (9 kDa antigen) |
| 114 | YP_001703235.1 | Hypothetical protein MAB_2500 |

| | Protein name | Description |
|---|---|---|
| 115 | YP_001703267.1 | Hypothetical protein MAB_2532 |
| 116 | YP_001703294.1 | Probable peptidyl-prolyl cis-trans isomerase |
| 117 | YP_001703315.1 | Hypothetical protein MAB_2580c |
| 118 | YP_001703426.1 | Conserved hypothetical protein |
| 119 | YP_001703431.1 | Hypothetical protein MAB_2697c |
| 120 | YP_001703460.1 | Hypothetical invasion protein Inv2 |
| 121 | YP_001703461.1 | Hypothetical invasion protein Inv1 |
| 122 | YP_001703472.1 | Probable thioredoxin TrxB |
| 123 | YP_001703474.1 | Conserved hypothetical protein |
| 124 | YP_001703532.1 | Conserved hypothetical protein |
| 125 | YP_001703533.1 | Conserved hypothetical protein |
| 126 | YP_001703534.1 | Conserved hypothetical protein |
| 127 | YP_001703538.1 | Conserved hypothetical protein |
| 128 | YP_001703539.1 | Lipoprotein LprG precursor (27 kDa lipoprotein) |
| 129 | YP_001703585.1 | Hypothetical protein MAB_2852c |
| 130 | YP_001703608.1 | Beta-lactamase precursor (Penicillinase) |
| 131 | YP_001703611.1 | Conserved hypothetical protein |
| 132 | YP_001703677.1 | Putative Mce family protein |
| 133 | YP_001703703.1 | Putative oxidoreductase |
| 134 | YP_001703704.1 | Conserved hypothetical protein |
| 135 | YP_001703713.1 | Putative lipoprotein LppU |
| 136 | YP_001703741.1 | Probable RNA polymerase sigma factor RpoD (Sigma-A) |
| 137 | YP_001703763.1 | Conserved hypothetical protein |
| 138 | YP_001703783.1 | Putative glutamate ABC transporter, periplasmic protein |
| 139 | YP_001703834.1 | Putative short chain dehydrogenase/reductase |
| 140 | YP_001703883.1 | Hypothetical protein MAB_3152c |
| 141 | YP_001703896.1 | Conserved hypothetical protein |
| 142 | YP_001703912.1 | Probable lipoprotein LppI |
| 143 | YP_001703950.1 | Hypothetical protein MAB_3220 |
| 144 | YP_001703991.1 | Probable lipoprotein LpqH precursor |
| 145 | YP_001703994.1 | Hypothetical protein MAB_3264c |
| 146 | YP_001704077.1 | Conserved hypothetical protein |
| 147 | YP_001704078.1 | Conserved hypothetical protein |
| 148 | YP_001704079.1 | Conserved hypothetical protein |
| 149 | YP_001704120.1 | Probable FeIII-dicitrate-binding periplasmic lipoprotein |
| 150 | YP_001704184.1 | Conserved hypothetical protein |
| 151 | YP_001704212.1 | NADPH-ferredoxin reductase FprA |
| 152 | YP_001704214.1 | Conserved hypothetical protein |
| 153 | YP_001704392.1 | Conserved hypothetical protein |
| 154 | YP_001704455.1 | Hypothetical protein MAB_3727c |
| 155 | YP_001704460.1 | 10 kDa chaperonin (GroES) |
| 156 | YP_001704474.1 | Conserved hypothetical protein |
| 157 | YP_001704481.1 | Conserved hypothetical protein |
| 158 | YP_001704491.1 | Probable cutinase cut2 precursor |
| 159 | YP_001704493.1 | Probable cutinase cut3 precursor |
| 160 | YP_001704494.1 | Probable cutinase cut3 precursor |
| 161 | YP_001704528.1 | Conserved hypothetical protein |
| 162 | YP_001704538.1 | Probable cutinase Cut4 |
| 163 | YP_001704581.1 | Conserved hypothetical protein |
| 164 | YP_001704582.1 | Conserved hypothetical protein |
| 165 | YP_001704583.1 | Putative lipoprotein LprC |
| 166 | YP_001704584.1 | Putative lipoprotein LprB precursor |
| 167 | YP_001704644.1 | Probable conserved secreted protein |
| 168 | YP_001704647.1 | Putative short chain dehydrogenase/reductase |
| 169 | YP_001704700.1 | Conserved hypothetical protein |
| 170 | YP_001704704.1 | Possible thioredoxin |
| 171 | YP_001704759.1 | Putative Mce family protein |

| | Protein name | Description |
|---|---|---|
| 172 | YP_001704761.1 | Putative Mce family protein |
| 173 | YP_001704762.1 | Putative Mce family protein |
| 174 | YP_001704763.1 | Putative Mce family protein |
| 175 | YP_001704781.1 | Putative short chain dehydrogenase/reductase |
| 176 | YP_001704789.1 | Conserved hypothetical protein |
| 177 | YP_001704801.1 | Lipoprotein LpqH precursor |
| 178 | YP_001704807.1 | Conserved hypothetical protein |
| 179 | YP_001704841.1 | Conserved hypothetical protein |
| 180 | YP_001704876.1 | Hypothetical MCE-family protein LprN |
| 181 | YP_001704911.1 | Superoxide dismutase |
| 182 | YP_001704950.1 | Probable glutamine-binding protein GlnH |
| 183 | YP_001704963.1 | Putative amino acid ABC transporter, substrate-binding protein |
| 184 | YP_001705003.1 | Probable conserved lipoprotein DsbF |
| 185 | YP_001705011.1 | Hypothetical protein MAB_4284c |
| 186 | YP_001705017.1 | Conserved hypothetical protein |
| 187 | YP_001705018.1 | Conserved hypothetical protein |
| 188 | YP_001705019.1 | Putative transcriptional regulator, LuxR family |
| 189 | YP_001705025.1 | Conserved hypothetical protein |
| 190 | YP_001705026.1 | Conserved hypothetical protein |
| 191 | YP_001705043.1 | Conserved hypothetical protein |
| 192 | YP_001705051.1 | Hypothetical protein MAB_4325c |
| 193 | YP_001705115.1 | Putative ABC transporter, periplasmic substrate-binding |
| 194 | YP_001705128.1 | Hypothetical protein MAB_4404 |
| 195 | YP_001705129.1 | Putative serine esterase, cutinase family |
| 196 | YP_001705181.1 | Putative secreted hydrolase |
| 197 | YP_001705235.1 | Putative Mce family protein |
| 198 | YP_001705236.1 | Putative Mce family protein |
| 199 | YP_001705238.1 | Putative Mce family protein |
| 200 | YP_001705251.1 | Conserved hypothetical protein |
| 201 | YP_001705260.1 | Conserved hypothetical protein |
| 202 | YP_001705287.1 | Putative Mce family protein |
| 203 | YP_001705290.1 | Putative Mce family protein |
| 204 | YP_001705318.1 | Putative Mce family protein |
| 205 | YP_001705321.1 | Putative Mce family protein |
| 206 | YP_001705329.1 | Conserved hypothetical protein |
| 207 | YP_001705350.1 | Hypothetical protein MAB_4627 |
| 208 | YP_001705424.1 | Conserved hypothetical protein |
| 209 | YP_001705462.1 | Possible beta-1,3-glucanase |
| 210 | YP_001705496.1 | Hypothetical protein MAB_4774c |
| 211 | YP_001705499.1 | Possible cellulase CelA (endoglucanase) |
| 212 | YP_001705504.1 | Conserved hypothetical protein |
| 213 | YP_001705523.1 | Possible twin-arginine translocation pathway |
| 214 | YP_001705574.1 | Phosphate ABC transporter, periplasmic protein |
| 215 | YP_001705620.1 | Single-stranded DNA-binding protein |
| 216 | YP_001705625.1 | Hypothetical protein MAB_4903 |
| 217 | YP_001705636.1 | Hypothetical protein MAB_4914c |
| 218 | YP_001705644.1 | Putative short-chain dehydrogenase/reductase |
| 219 | YP_001705646.1 | Conserved hypothetical protein |
| 220 | YP_001705647.1 | Hypothetical protein MAB_4925 |
| 221 | YP_001705657.1 | MutT/NUDIX family protein |
| 222 | YP_001705663.1 | Thioredoxin (Trx) |
